# Effect of integrated health system leading-managing-and-governing for results model on institutional delivery: Team-based quasi-experiment

**DOI:** 10.1101/775858

**Authors:** Yeshambel Agumas Ambelie, Getu Degu Alene, Damen Hailemariam Gebrekiros

## Abstract

**Objective:** The objective of this study is to examine, based on theory of change, whether integrated leading-managing-and-governing for results model is plausible cause of improved institutional delivery.

**Methods:** A team-based quasi-experimental study was conducted. One-hundred-thirty-four health facility teams were enrolled in the study. Teams were allocated to intervention and control groups in a 1:1 ratio, non-randomly. End line institutional delivery was the dependent variable while the group (main predictor) and the baseline institutional delivery (covariate) were independent variables. The intervention that was given over six months was integrated leading-managing-and-governing for results model. The institutional deliveries were measured with percentages whilst the group was measured with exposure status (yes or no) to the intervention. Data, from both groups, were collected at baseline and end line. Data were analyzed using analysis of covariance. Statistical significance was determined at (p<.05). The main effect of the intervention was determined by 95% CI, presented in the contrast results.

**Results:** The adjusted mean institutional deliveries with 95% CI were 47.4 (46.2, 48.6) and 33.4 (32.2, 34.6) in the intervention and control groups, respectively. Contrast results showed that having an intervention group, p = .000, 95% CI (12.2, 15.8), of integrated leading-managing-and governing for results model significantly increased mean institutional delivery compared to having a control group.

**Conclusions:** This study provides some guidance regarding the plausible causation of integrated leading-managing-and-governing for results model on institutional delivery. It would serve as a baseline in identifying true causation using a randomized design.

## Introduction

Strong health system is required to address global concerns such as Universal Health Coverage (UHC)[1,2]. To realize this, six critical health system building blocks are identified[3,4]. This includes service delivery, health workforce, medical products, health information systems, healthcare financing, and leadership and governance. But, leadership and governance is remained the most challenging to measure, particularly in Low and Middle-Income Country (LMIC) health systems[5,6]. Perhaps, it might be due to lack of scientifically reliable and empirically scalable practices.

Despite this challenge, Integrated Leading-Managing-and-Governing (ILMG) for results model (**Fig 1**), centered in the leadership development program, has been employed over 50 LMIC health systems including Ethiopia[5,6]. Mainly, this has been implemented as a pilot project through the technical support and budget aid from international organizations like Management Science for Health (MSH), John Snow Inc. (JSI) and United States Agency for International Development (USAID)[3,7].

**Fig 1.**
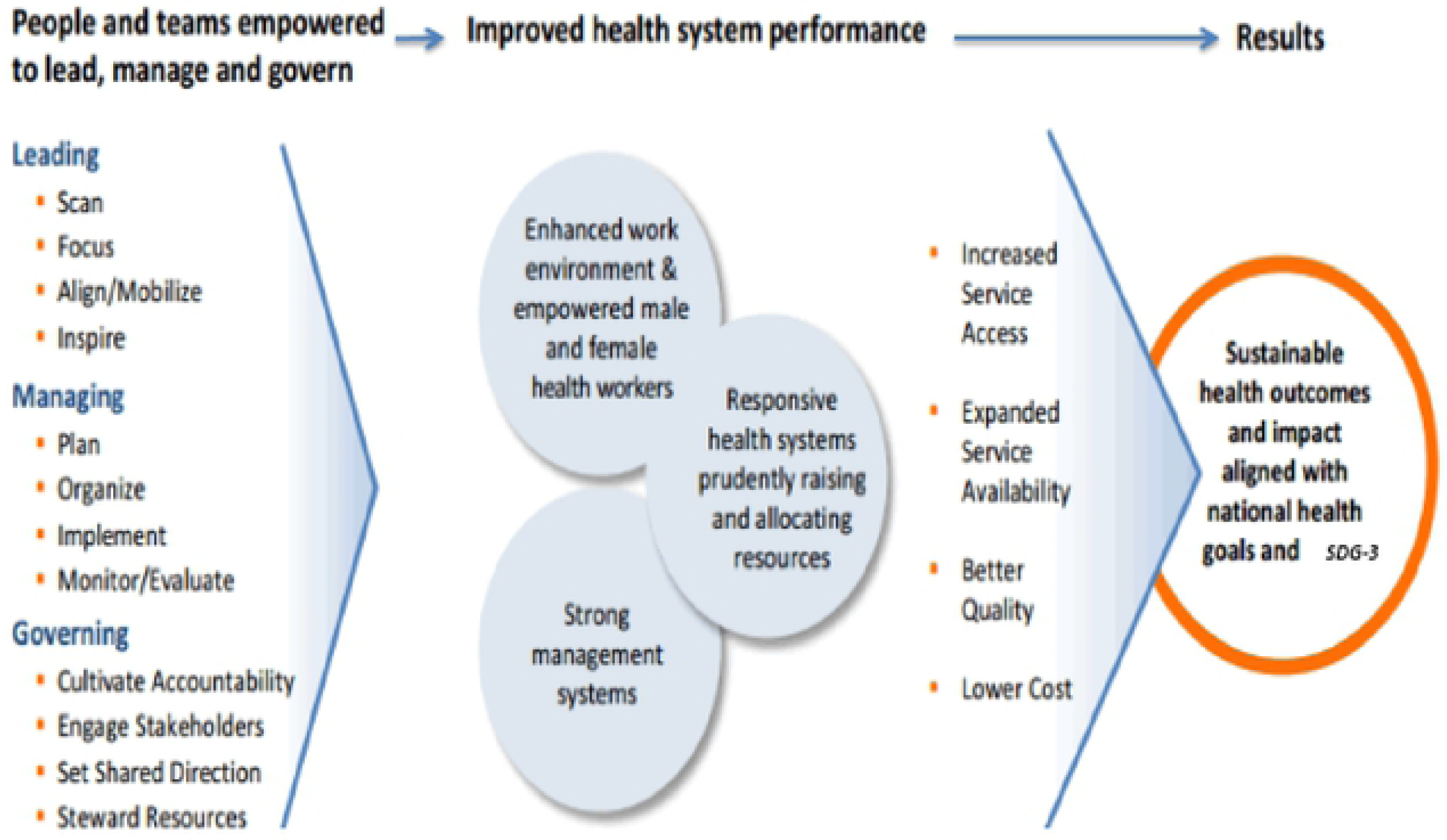
Conceptual model: ILMG for results (Source: MSH, 2015)

In fact, in the beginning, the model held only integrated leading and managing practices[7-9]. But, a decade later, it holds the current structure (**Fig 1**) through incorporating governing practices[3,10]. Additionally, using factor analysis technique, the current authors reported four integrated latent factors of the three paths[10]. These are compliance with principles, strategic sensitivity, system building and contextual thoughtfulness. Such findings strengthen the challenging characteristics of measuring the leadership and governance building block. The model (**Fig 1**), particularly in Ethiopia, is applied in the USAID transform primary health care project health facilities, with a goal of ending preventable maternal mortality. Away from its enormous expansion, only limited studies report the effect of the model on improved health system performance and sustained health outcomes[6,8,9,11]. The studies done in Kenya, Egypt and Mozambique reported that applying the model increased 10%, 41% and 10% average coverage rate on selected health-service delivery indicators. However, the latter two studies were done retrospectively for evaluating pilot projects[9,11].

On the contrary, the study done in Afghanistan reported that there was no statistically significant effect of the intervention on health system performance[12]. Rather, it showed that many indicators worsened in the intervention group.

Generally, the previous studies lack either using control group[9,11] or controlling plausible confounding factors[8,9,11,12].

Thus, researching the effect of any initiative led by Theory of Change (ToC) and using rigorous methodology would be important in generating better evidences[13,14]. ToC refers a systematic and cumulative study of the links between input, activities, output, outcome and context of the initiative[14]. There are three identified attributes to achieve the potential of ToC on initiatives: plausibility, doablity and testability [13,15]. Plausibility refers whether activities implemented should lead to desired outcomes; doablity is about availability of all resources to carry out the initiative, and testability explains presence of specific and complete ToC to track its progress in credible and useful ways. Moreover, the research done in Kenya acknowledged that using research outcome of interests that varied from team to team lead the analysis to focus on average coverage or service volume rather than on specific indicators[8]. To avoid this, they recommended focusing on either teams addressing the same indicator or a set of related indicators.

Therefore, this study aims at examining the effect of the ILMG for results model on Institutional Delivery (ID) using a prospective pre–post intervention no-treatment control group team-based[16] quasi-experimental study. Quasi-experiment is an empirical study design used to estimate the plausible causal impact of an intervention on its target population without random assignment[17,18]. The findings of this study would support evidence-based leveraging of the model at all levels that is either scale-up or re-design it; as well as serve as a baseline for future research.

## Methods

### Study design and teams

A prospective pre–post intervention no-treatment control group team-based quasi-experimental study was conducted. One-hundred-thirty-four health facility teams were enrolled in the study. These teams were allocated to intervention and control groups in a 1:1 ratio, non-randomly. Integrated leading-managing-and-governing for results model was given to the intervention group. Yet, the control group was followed without any intervention. Moreover, teams were intact and worked together over the intervention period.

### Intervention

The ILMG for results model was delivered over six-month period. Based on the intervention protocol (**S1 appendix**), basic concepts that enable the teams to face challenges, and achieve results were transferred with two consecutive off-site three-day workshops.

During the first workshop, the main task that the teams carried out was developing six-month project on ID using a tool called the Challenge Model (CM)[3,9] (***Fig 2***).

**Fig 2.**
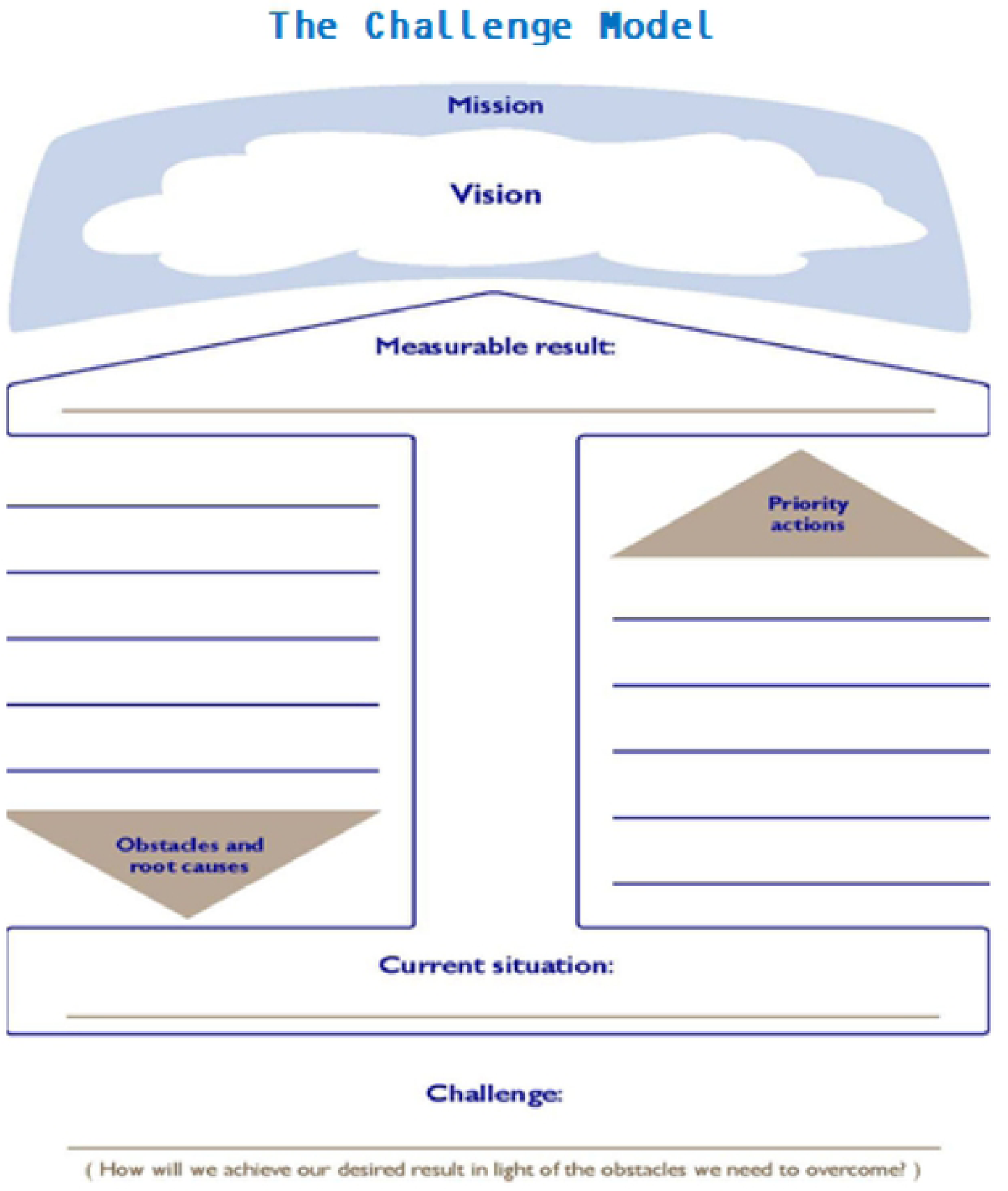
The challenge Model (Source: Mansour M et al, 2010)

The activities, elements of the CM, that the teams worked step-by-step were: reviewed their respective facility mission; set a shared vision in lined with facility vision; developed six-month Measurable Result (MR); assembled current situation (baseline); identified obstacles and root causes[19-21]; developed inspirational challenge statement by combining the MR and obstacles; and designed priority actions to avert obstacles. Moreover, they identified potential stakeholders to align and mobilize resources for better result.

With all the above activities done, the teams were sent back to their respective working place, taking an assignment of sharing and validating the project with the other staff and key stakeholders. Additionally, teams were encouraged to exercise the ILMG for results model. After average period of one-month, the teams were called back for the second workshop. It was began with presentations and discussion on the validated projects. Furthermore, teams were facilitated to develop action plan and monitoring and evaluation (M&E) plan using respective planning formats (**S2 and S3 Appendixes**). Moreover, concepts of coaching using Observe, Ask, Listen, Feedback and Agree (OALFA) technique; communication using effective model (**S4 Appendix**); managing facility resource and health services delivery were presented and discussed. By the end of the third day, the teams were sent back to their respective working place for the actual implementation of their projects.

Another-month after, based-on the OALFA technique, the facilitators including the investigators did on-site coaching visit for each team. Facilitators were certified experts for integrated leadership-management-and governance Trainer of Trainees (TOT), from the Ethiopian federal ministry of health. Participatory, enquiry-based and practice-oriented facilitation approaches were employed. In addition, brainstorming ideas, insight-invoking questions, role-plays, group discussion, case studies and work place assignments were also used. Moreover, concise and comprehensive notes, tables and figures were distributed to teams as needed.

### Variables and measuremnts

The Dependent Variable (DV) was the end line ID while the main independent variable was the group. Another independent variable, the baseline ID that had an influence on the DV was considered as a covariate. Regarding to measurements, both the baseline and end line institutional deliveries were measured with percentage means. The groups were measured with exposure status (yes or no) to the intervention.

### Data collection and analysis

The baseline and end line data were collected from teams of each group. Before getting to the final analysis, five stages of data analyses were conducted. These were done using the statistical package for the social sciences version 20. First, descriptive analysis was done to characterize the ordinary mean ID. Second, assumptions of no presence of significant outliers and approximately normally distributed data for each group were assessed by boxplot and Shapiro-Wilk test.

Third, in the absence of the covariate, the effect of the group on the DV was tested using Analysis of Variance (ANOVA). The output indicated significant result: F (1,132) = 79.0, partial eta squared = .37, and p < .001. Partial eta squared measured the proportion of the total variance (effect size) on the DV that is associated with the membership of different groups defined by a group[22]. For example, the above output showed that the group (intervention and control) accounted for 37% of the variability on the DV.

Fourth, group-covariate interaction effect was checked using custom model of Analysis of Covariance (ANCOVA). It tested differences between group means when we knew that an extraneous variable affected the outcome of interest[23,24]. Observing at the p-value of the analysis output: F (1,130) = 1.6, partial eta squared = .01 and p = .21, it was obvious that the covariate was not significantly predicted the DV. The other important output displayed from this analysis was the result of Levene’s test: F (1, 132) = 58.5 and p = 000. It indicated that the group variances were not equal and hence the assumption of homogeneity of variance was violated. This further showed that we failed to reject the null hypothesis in that there was no group by covariate effect on DV. Alternatively, the covariate had the same correlation with the DV for both intervention and control groups; and the correlation between the covariate and DV was while they differ for intervention and control groups. Precisely speaking, there was no Lord’s paradox[24-26]. Fifth, the effect of the group on the covariate was also tested using ANOVA. The output presented that the group was not significantly predicted the covariate: F (1,132) = 2.8, partial eta squared = .02, and p = .09.

Considered the above outlooks, ANCOVA with full factorial model was conducted to evaluate the main effect of the group on DV. The 95% CI from the contrast results was used in determining the main effect of the intervention. From this output, two things were considered: (1) did significant value less than .05, and (2) did not the CI include zero.

The CI here was the difference between means, the original means adjusted for the covariate that showed the likely value in the population. In reality, if the difference between means is zero, then it tells there is no difference between the groups. If the CI does not contain zero, it means that the effect in population is likely to be bigger or smaller than zero.

### Ethical considerations

The current study was registered at clinical trials.gov with identifier NCT03639961. Additionally, ethical clearance was secured from Bahir Dar University (BDU) with a protocol record 090/18-04. Moreover, written consent was obtained from each members of study teams; and data were protected.

## Results

### Ordinary means

Table 1 displays the ordinary mean and standard deviation (SD) of the baseline and end line ID with 95% CI. The mean difference between the baseline and end line ID was 14.6±7.2 in the intervention group, whereas, it was 1.1±2.2 in the control group.

**Table 1.**
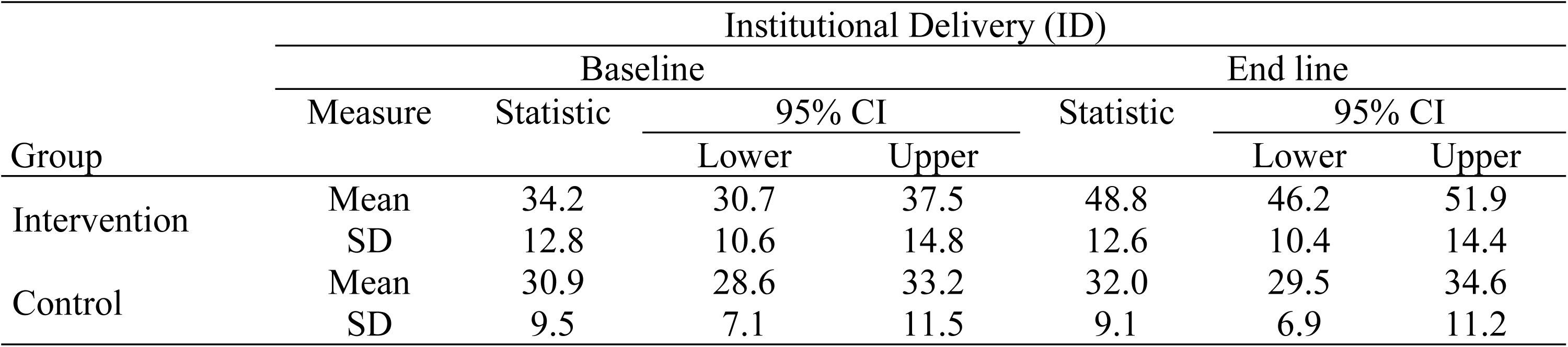
Ordinary mean (SD) baseline and end line ID (n= 134)

### Estimated means

Table 2 presents the adjusted mean end line ID for both groups that were the original means adjusted for the covariate. The mean values had changed compared to those found in the ordinary mean (**Table 1**).

**Table 2.** Adjusted mean end line ID (n= 134)

| Group | Mean | Std. Error | 95% Confidence Interval |  |
| --- | --- | --- | --- | --- |
|  |  |  | Lower Bound | Upper Bound |
| Intervention | 47.4 | .627 | 46.2 | 48.6 |
| Control | 33.4 | .627 | 32.2 | 34.6 |

*Note: The covariate appearing in the model was evaluated at the following value: baseline ID = 32.6.*

### The main effect of the group on the DV

Table 3 informs that there was an overall statistically significant difference on the DV between the groups once their means had been adjusted for the covariate. As highlighted in the table, there was statistically significant difference between adjusted means: F (1,131) = 247.2, partial eta squared =.65 and P<.001. Considered the partial eta squared value, the main effect size of the group on DV was 65%. This showed that including the covariate increased the group’s effect size on the DV from 37% (explained in methods part) to 65%.

**Table 3.** Outputs of between-subjects effects on end line ID, ANCOVA (n = 134)

| Source | Type III Sum of Squares | Df | Mean Square | F | Sig. | Partial Eta Squared |
| --- | --- | --- | --- | --- | --- | --- |
| Corrected Model | 21955.8 | 2 | 10977.9 | 421.7 | .000 | .87 |
| Intercept | 2169.6 | 1 | 2169.6 | 83.3 | .000 | .39 |
| Baseline ID | 12460.4 | 1 | 12460.4 | 478.6 | .000 | .79 |
| Group | 6435.3 | 1 | 6435.3 | 247.2 | .000 | .65 |
| Error | 3410.4 | 131 | 26.0 |  |  |  |
| Total | 244108.0 | 134 |  |  |  |  |
| Corrected Total | 25366.2 | 133 |  |  |  |  |

*Note: R Squared = .87, Adjusted R Squared = .86*

Yet, Table 3 also displayed that the covariate had significant effect at (P<.001). Thus, to interpret such outputs, double testing using contrast results (K matrix) (**Table 4**) was used.

The output indicated significant result (P<.001), and the 95% CI was (12.2, 15.8). This showed that the main effect of the intervention was somewhere between this CI.

**Table 4.**
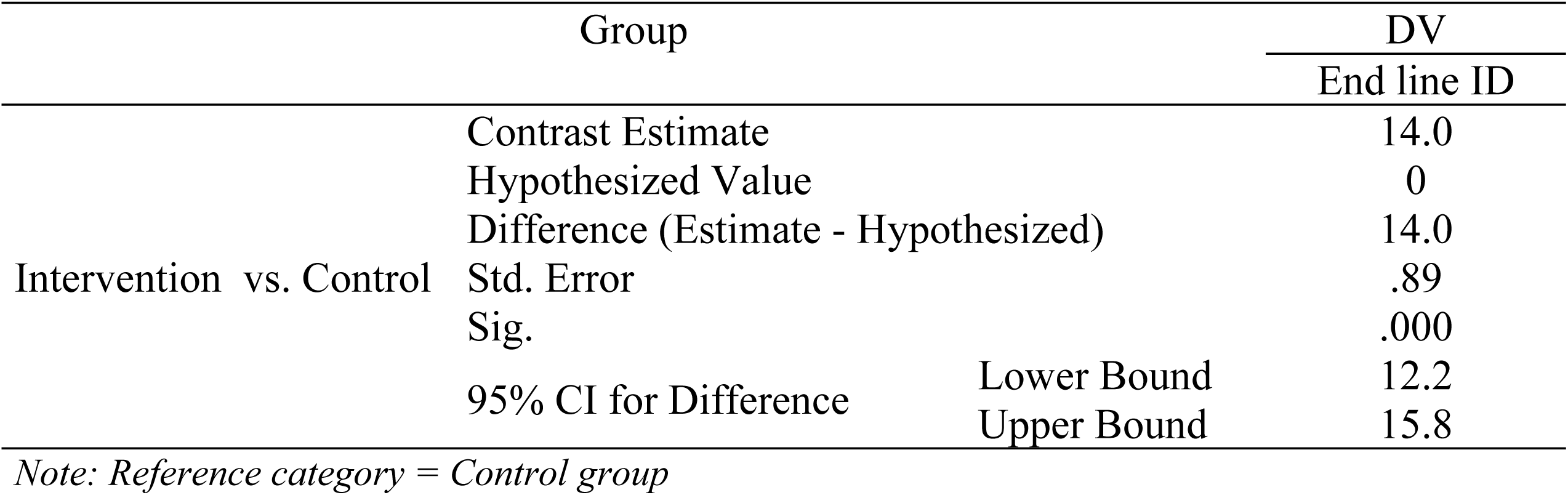
K Matrix

## Discussion

The current study findings inform that the ILMG for results model intervention causes statistically significant difference on mean ID between the groups. This plausible causation is supported by a study done in Kenya[8]. Differently, the current study shows the effect size by adjusting the original means for the covariate. This has three-fold purposes[18]. First, it reduces within-group error variance that is the intervention effect bias or specification error. Second, it eliminates potential confounders since there is no preexisted group differences systematically on more than it is. Last, it provides additional evidence of causality.

The current finding is also supported by other previous studies[9,11]. However, unlike these studies, the current study controlled a plausible covariate (noted earlier) and used control group. Using control group helps to identify assumption attributes in trends between the groups that occur at the same time as the intervention to that intervention.

The other distinction of the current study is that it used the model that integrated leading-managing-and-governing practices (Fig 1) while the other studies used either leading and managing practices[8,9] or governing practices[12]. Interestingly, the effect of balancing and integrating leading managing and governing practices in improving service-delivery outcome in a turbulent environment is similar to keeping the seat of a three-legged stool horizontal while sitting on rough ground[27] (**Fig 3**).

**Fig 3.**
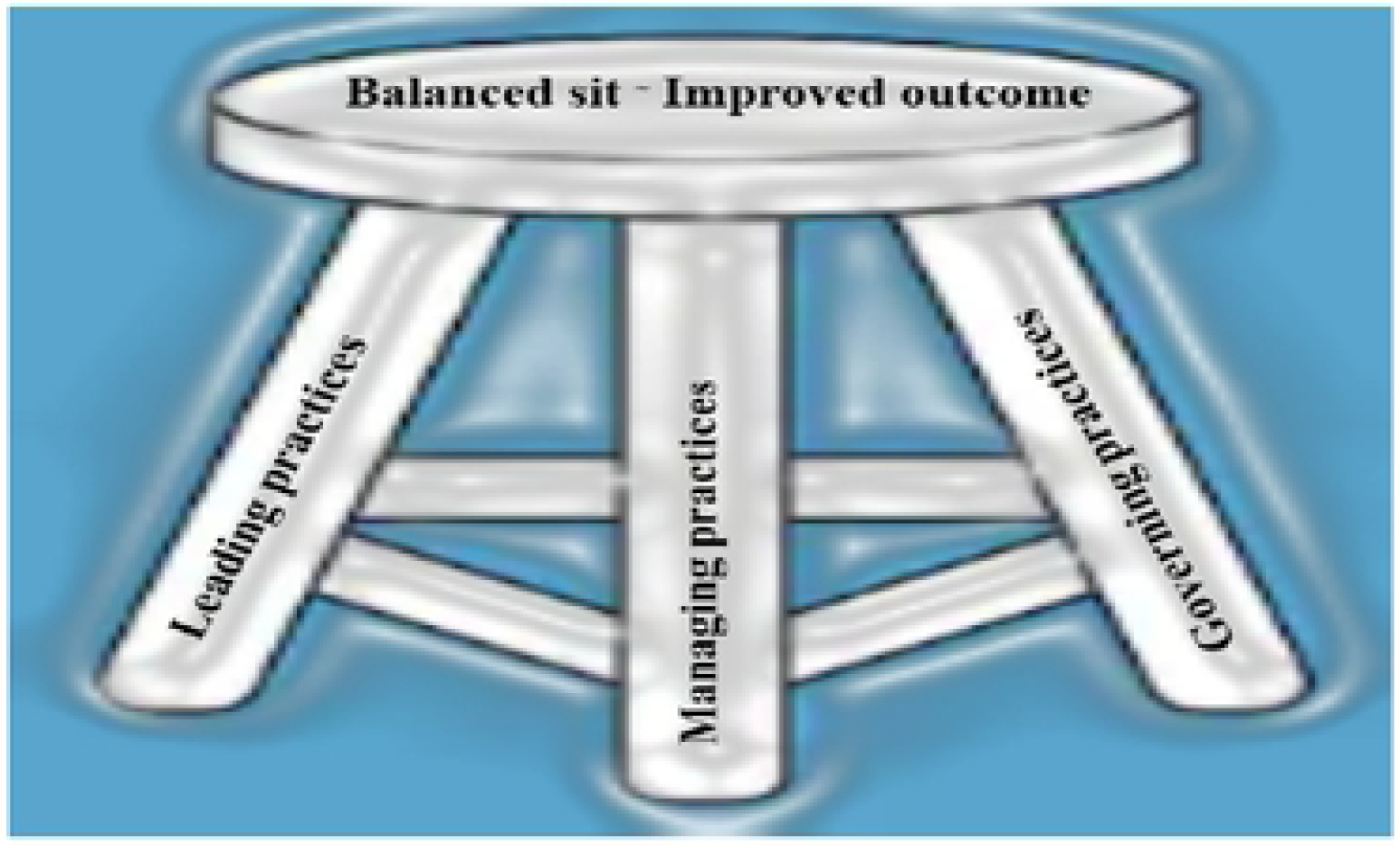
Illustration of a sit on adjusted three-legged stool with effect of ILMG practices on services outcomes.

On the contrary, the study done in Afghanistan reported that there was no statistically significant effect of the intervention on health system performance or health outcomes[12]. Rather, it showed that many indicators worsened in the intervention group. As explained by the authors of the study, the intervention environment was fragile and conflict affected in the study period. This supports the significant influence of the turbulent environment to achieve significant results through interventions.

In the current study, the adjusted mean ID (**Table 2**) compared with the ordinary mean ID (**Table 1**) is less in the intervention group, but greater in the control group. This implies that adjusting the mean by removing error variance in the DV that associates with the covariate provides unbiased or uncontaminated mean.

The current adjusted means in both groups are also greater compared with the 2016 Ethiopian demographic health survey ID report (26%). However, compared with the 2019 demographic health survey ID report (48%): the current adjusted mean ID in the intervention group is similar, but it is less in the control group. These mean institutional deliveries are also by far lower than the target (90%) indicated in the national health sector plan, 2015-2019. Taking into consideration, perhaps, the national survey result includes data from big cities, evidence-based investment on ILMG for results model ought to be important.

In spite of the above implications, there were potential limitations in conducting this study. The first limitation identified was non-randomization that is the major weakness of quasi-experimental design. This weakness brought another challenge that was whether ANCOVA is used in alike data. Yet, two dimensions support its application. First, if the group have caused the difference on covariate beyond randomization[28]. Second, if the authors are certain that the group could not have affected the covariate[29]. Since there was no preexisted group that affected the covariate in the current study, ANCOVA was applied. In fact, this analysis technique is developed to increase the power of the test of the predictor variable[23,24]. It does this through removing error variance in the DV that is associated with the covariate[24]. The second important threat to establishing causality was the statistical principle of regression to the mean[18]. This widespread statistical phenomenon can result in wrongly concluding that an effect is due to the intervention when in reality it is due to chance. Here, though, the degree of caution was diminished by implementing the intervention on the real-world setting, limiting generalizability of results is unavoidable[5].

The third potential limitation was the short duration of the intervention. Six months may not be enough time to overcome barriers and achieve significant result. Nevertheless, if it was more than this with similar study design that lacks isolation and temporal precedence, contamination will be a threat on the other way round. Even with this time, though no team was recorded as loss to follow up, around 11% of intervention teams reported that at least one team member transferred to a new area at the time of intervention.

The last challenge, to the best of our knowledge, was dearth of available literatures on testing ILMG for results model, which of course limited the depth of our discussion.

## Conclusions

This study provides some guidance regarding the plausible causation of integrated leading-managing-and-governing for results model on institutional delivery. It would support evidence-based-leveraging of the model in similar settings. It would also serve as a baseline for future research, possibly, considering randomization to identify true causation.

## Acknowledgements

Our sincere appreciation go to the study participants, intervention facilitators, data collectors, and data supervisors, for their valuable contribution.

## Supporting information

**S1 Appendix.** Intervention protocol

**S2 Appendix.** Action plan planning format

**S3 Appendix.** M&E plan planning format

**S4 Appendix.** Effective communication model

## References

1. Murray CJ, Frenk J (2000) A framework for assessing the performance of health systems. Bulletin of the world Health Organization 78: 717–731.

2. Peters DH, El-Saharty S, Siadat B, Janovsky K, Vujicic M (2009) Improving health service delivery in developing countries: from evidence to action: The World Bank.

3. Management Sciences for Health. Rice JA, Shukla, Mahesh, Johnson Lassner, Karen et al. (2015) Leaders Who Govern. June 2015.

4. Bhutta ZA, Chopra M, Axelson H, Berman P, Boerma T, et al. (2010) Countdown to 2015 decade report (2000–10): taking stock of maternal, newborn, and child survival. The lancet 375: 2032–2044.

5. Rauscher M, Walkowiak H, Djara MB (2018) Leadership, Management, and Governance Evidence Compendium.

6. Daire J, Gilson L, Cleary S (2014) Developing leadership and management competencies in low and middle-income country health systems: a review of the literature. Cape Town: Resilient and Responsive Health Systems (RESYST).

7. Galer JB, Vriesendorp S, Ellis A (2005) Managers who lead: a handbook for improving health services.

8. La Rue KS, Alegre JC, Murei L, Bragar J, Thatte N, et al. (2012) Strengthening management and leadership practices to increase health-service delivery in Kenya: an evidence-based approach. Human resources for health 10: 25.

9. Mansour M, Mansour JB, El Swesy AH (2010) Scaling up proven public health interventions through a locally owned and sustained leadership development programme in rural Upper Egypt. Human Resources for Health 8: 1.

10. Yeshambel AA, Getu DA, Damen HG (21 June 2019 PREPRINT (Version 1)) Performance capacity of the health system workforce towards integrated leading-managing-and-governing practices in Ethiopia: with special reference to the application of factor analysis and ordinal logistic regression.

11. Perry C (2008) Empowering primary care workers to improve health services: results from Mozambique’s leadership and management development program. Human resources for health 6: 14.

12. Anwari Z, Shukla M, Maseed BA, Wardak GFM, Sardar S, et al. (2015) Implementing people-centred health systems governance in 3 provinces and 11 districts of Afghanistan: a case study. Conflict and health 9: 2.

13. Connell JP (1995) New Approaches to Evaluating Community Initiatives. Concepts, Methods, and Contexts. Roundtable on Comperhensive Community Initiatives for Children and Families: ERIC.

14. Connell JP, Kubisch AC (1998) Applying a theory of change approach to the evaluation of comprehensive community initiatives: progress, prospects, and problems. New approaches to evaluating community initiatives 2: 1–16.

15. Brown P (1995) The role of the evaluator in comprehensive community initiatives. See Ref 24a: 201–225.

16. Salas E, DiazGranados D, Klein C, Burke CS, Stagl KC, et al. (2008) Does team training improve team performance? A meta-analysis. Human factors 50: 903–933.

17. DiNardo J (2010) Natural experiments and quasi-natural experiments. Microeconometrics: Springer. pp. 139–153.

18. Harris AD, McGregor JC, Perencevich EN, Furuno JP, Zhu J, et al. (2006) The use and interpretation of quasi-experimental studies in medical informatics. Journal of the American Medical Informatics Association 13: 16–23.

19. Ilie G, Ciocoiu CN (2010) Application of fishbone diagram to determine the risk of an event with multiple causes. Management Research and Practice 2: 1–20.

20. Ballantyne D (1990) Coming to grips with service intangibles using quality management techniques. Marketing Intelligence & Planning 8: 4–10.

21. Collins JC, Porras JI (1996) Building your company’s vision. Harvard business review 74: 65–&.

22. Richardson JT (2011) Eta squared and partial eta squared as measures of effect size in educational research. Educational Research Review 6: 135–147.

23. Cochran WG (1957) Analysis of covariance: its nature and uses. Biometrics 13: 261–281.

24. Miller GA, Chapman JP (2001) Misunderstanding analysis of covariance. Journal of abnormal psychology 110: 40.

25. Jin P (1992) Toward a reconceptualization of the law of initial value. Psychological Bulletin 111: 176.

26. Lord HW, Shulman Y (1967) A generalized dynamical theory of thermoelasticity. Journal of the Mechanics and Physics of Solids 15: 299–309.

27. Levey S, Vaughn T, Koepke M, Moore D, Lehrman W, et al. (2007) Hospital leadership and quality improvement: rhetoric versus reality. Journal of Patient Safety 3: 9–15.

28. Overall JE, Woodward JA (1977) Nonrandom assignment and the analysis of covariance. Psychological Bulletin 84: 588.

29. Wildt AR, Wildt AA, Ahtola O, Ahtola OT (1978) Analysis of covariance: Sage.

